# Newly developed single-cell computational approach elucidates the stabilization of *Oct4* expression in the E3.25 mouse preimplantation embryo

**DOI:** 10.1101/476127

**Authors:** Daniela Gerovska, Marcos J. Arauzo-Bravo

## Abstract

The time of onset of the second cell fate decision in the mouse preimplantation embryo is still unknown. Ohnishi *et al.* (2014) looked for cell heterogeneity in the ICM at E3.25 that could indicate the time preceding the apparent segregation of PE and EPI at E3.5, but were not able to detect an early splitting transcriptomics event using state-of-the-art clustering techniques. We developed a new clustering algorithm, hierarchical optimal *k*-means (HO*k*M), and identified from single cell (sc) transcriptomics microarray data two groups of ICM cells during the 32 to 64 mouse embryo transition: from embryos with less than 34 cells, and more than 33 cells, corresponding to two developmental sub-stages. The genes defining these sub-stages indicate that the development of the embryo to 34 cells triggers a dramatic event as a result of which *Bsg* is high expressed, the canonical Wnt pathway is activated, *Oct4* is stabilized to high expression and the chromatin remodeling program is initialized to establish a very early narve pluripotent state from the preceding totipotency. We characterized our HO*k*M partition comparing with independent scRNA-seq datasets. It was staggering to discover that from the 3.4360×10^10^ possible bi-partitions of the E3.25 data of Ohnishi *et al.* (2014), our HO*k*M discovered one partition that shares the biological features of the early and late 32 ICM cells of Posfai *et al.* (2017). We propose that the stabilization of *Oct4* expression is a non-cell autonomous process that requires a minimal number of four inner cell contacts acquired during the transition from a homogeneous outer-cell environment to a heterogeneous inner/outer cell environment formed by the niche of a kernel of at least six inner cells covered by trophectoderm.

## Introduction

The mouse preimplantation development begins with the division of the 1-cell zygote to progressively smaller cells blastomeres, forming the morula, which at the 8-cell stage compacts. As a result of a first cell fate decision, the morula ball forms a hollow and becomes blastocyst with outer cells forming the trophectoderm (TE) and interior cells, the inner cell mass (ICM). The ICM sets aside from outer cells in two successive waves of asymmetric cell division, 8-16-cell and 16-32-cell transitions, with Morris *et al.* (2010) reporting an additional third one. A second cell fate decision involves the segregation of the ICM, appearing morphologically as a homogeneous cell population, into embryonic epiblast (EPI), pluripotent and progenitor of the future body, and primitive endoderm (PE), which becomes a morphologically distinct monolayer by implantation at E4.5 (Zernicka-Goetz *et al.*, 2009).

The mechanism governing the ICM cell specification in the early blastocyst is still unclear. Ohnishi *et al.* (2014) looked for cell heterogeneity in the ICM at E3.25 that could indicate early onset of the second cell fate decision, the time preceding the apparent segregation of PE and EPI at E3.5, but were not able to detect an early splitting transcriptomics event using state-of-the-art clustering techniques.

Oct4 is master regulator of pluripotency (Jaenisch and Young, 2008) and cornerstone not only in developmental biology but also in regenerative medicine for its role in somatic cellular reprogramming (Takahashi and Yamanaka, 2006; Kim *et al.*, 2009a; 2009b). *Oct4* is expressed *de novo* in the ICM after being down-regulated in the early cleavage stages (Rosner *et al.*, 1990), however the mechanism that reactivates *Oct4* in the ICM remains unknown.

To obtain a more complete picture of the cell specification events occurring between 32- and 64-cell stage, we developed a new clustering algorithm, with which we looked for structure in the heterogeneity during the 32-64 cell wave of divisions, for transcriptomics events explaining the loss of totipotency in the ICM, and for the mechanism behind the reactivation of *Oct4*. We reanalyzed the single single (sc) transcriptomics microarray data of Ohnishi *et al.* (2014) with the design of a new clustering algorithm, Hierarchical Optimal *k*-means (HO*k*M), and succeeded in identifying two groups of cells in the E3.25 ICM, one of cells from embryos with less than 34 cells, and another of cells from embryos with more than 33 cells, which we hypothesized characterized two developmental sub-stages in the mouse preimplantation embryo (named later E3.25-LNCs, less than 34 cells, and E3.25-HNC, more than 33 cells, respectively). Next, we calculated the differently expressed genes between the two groups, E3.25-LNCs and E3.25-HNCs, and thus we found *Oct4* among the top upregulated genes in the E3.25-HNCs. It is worth mentioning that the number of all possible partitions of the 36 single cell transcriptomics data set of E3.25 from Ohnishi *et al.* (2014) is the Bell number *B*_36_ = 3.8197×10^30^, and the number of all possible bi-partitions is the Stirling number of the second class ***S***_36,2_ = 3.4360×10^10^. It was staggering to discover that our HO*k*M partition was associated with a stabilization of *Oct4* at high expression level. Previously, Ohnishi *et al.* (2014) found bimodal expression of *Fgf4* within the E3.25 ICM cells, suggesting that as an early indication of future PE or EPI fate. We hypothesized that such bimodal expression of *Fgf4*, among others, could be traced to earlier developmental stages and performed a bifurcation analysis on the sc data of Ohnishi *et al.* (2014), spanning the time from E3.25, E3.5 to E4.5, while introducing an additional bifurcation point (between E3.25-LNC and E3.25-HNC) during E3.25 thanks to the newly identified E3.25 sub-stages. Thus we found that *Fgf4*, among other PE and EPI markers, has bimodal expression even before E3.25-LNC. To compare the transcriptional similarity of the E3.25-LNC and E3.25-HNC with other developmental stages, we calculated PCA and violin plots of the top upregulated genes in the E3.25-HNC, using the transcriptomics microarray data of Ohnishi *et al.* (2014) together with data from the same and the earlier developmental stages of the same Affymetrix platform from other authors. To check whether the E3.25-HNC are similar to either PE or EPI cells, we performed scatter plots between the E3.25-HNC and E3.5 PE or E3.5 EPI cells of Ohnishi *et al.* (2014). We used the availability of Fgf4-KO data in the dataset of Ohnishi *et al.* (2014) to compare the transcriptional similarity of the E3.25-LNC and E3.25-HNC with E3.25 Fgf4-KO, E3.5 Fgf4-KO and E4.5 Fgf4-KO. We have retrospectively validated our HO*k*M clustering algorithm on the E3.5 and E4.5 sc transcriptomics microarray data of Kurimoto *et al.* (2006) to group the PE and EPI cells, and since their sc microarray dataset is produced on the same platform, we used it together with the E3.5 and E4.5 data of Ohnishi *et al.* (2014) to find common and specific PE and EPI markers for E3.5 and E4.5. Then, we evaluated the closeness of all existent ESCs in the GEO base from the same Affymetrix platform to their *in vitro* ICM counterparts from Ohnishi *et al*.(2014). In order to validate the existence of developmental sub-stages around E3.25, we reanalyzed the scRNA-seq data from Posfai *et al.* (2017). Finally, we built a simple cell packaging model to explain the strong transcriptional changes between ICM cells from embryos with less than 34 cells and ICM cells from embryos with more than 33 cells.

## Results

### E3.25 ICM cells segregate into lower-(LNC) and higher-number-of-cell (HNC) clusters

The Principal Component Analysis (PCA) (Fig. 1) of transcriptomics data from oocyte till early post-implantation stages (Table 1) shows that first and second Principal Components reflect the embryo age and cell fate, respectively. Focusing on the E3.25-E3.5 stages using only the ICM single-cell data from Ohnishi *et al*. (2014) (Fig. 1B), revealed that E3.25 cells divide into two groups, one comprised mainly of cells from embryos with lower number of cells (LNCs), between 32 and 33, and cell 26 C41 IN from a 41-cell embryo, and another comprised of cells from embryos with higher number of cells (HNCs), between 34 and 50.

**Table 1.**
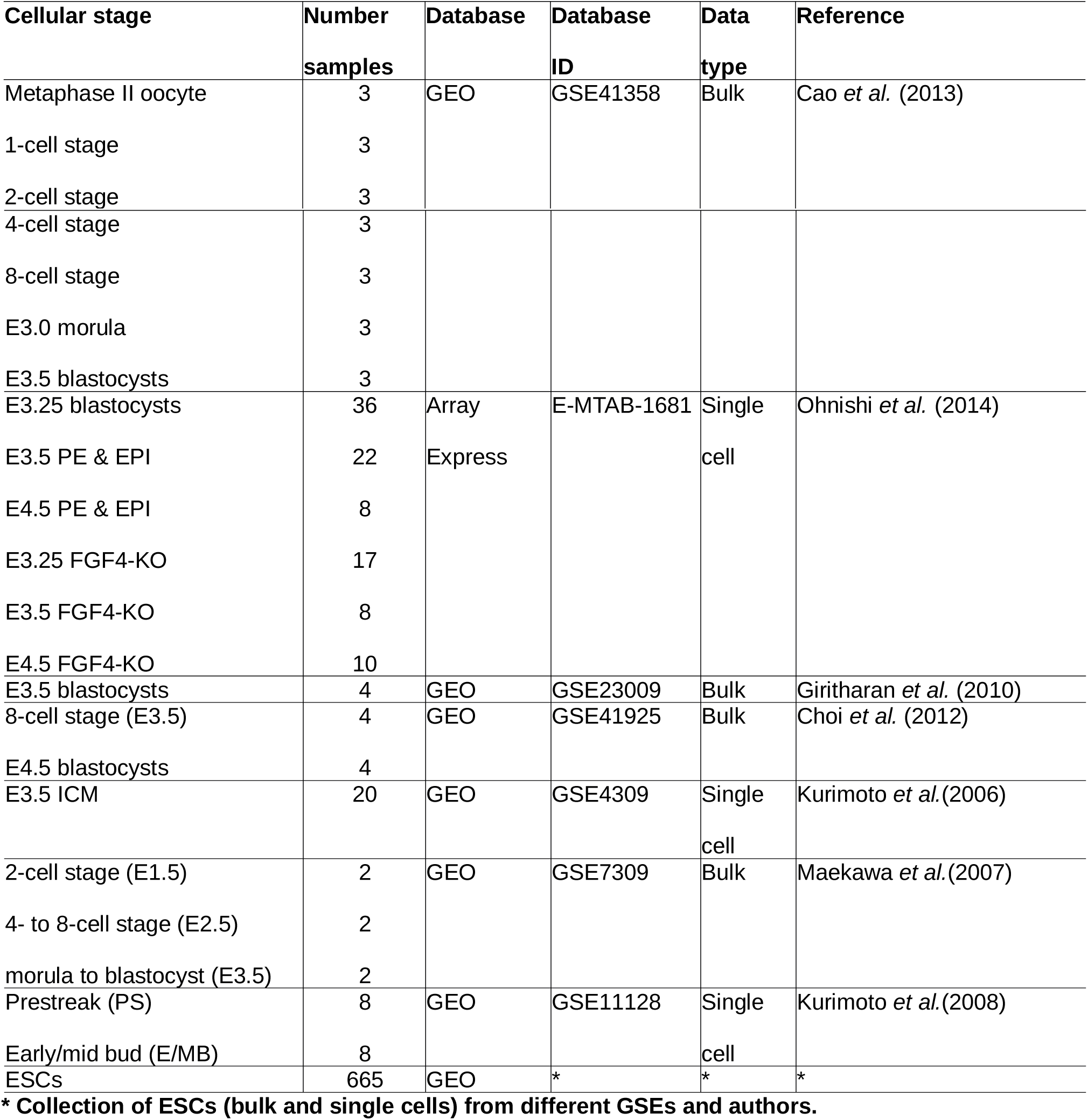
Datasets from the Affymetrix Mouse Genome 430 2.0 Array used in the different transcriptomics analyses.

**Figure 1.**
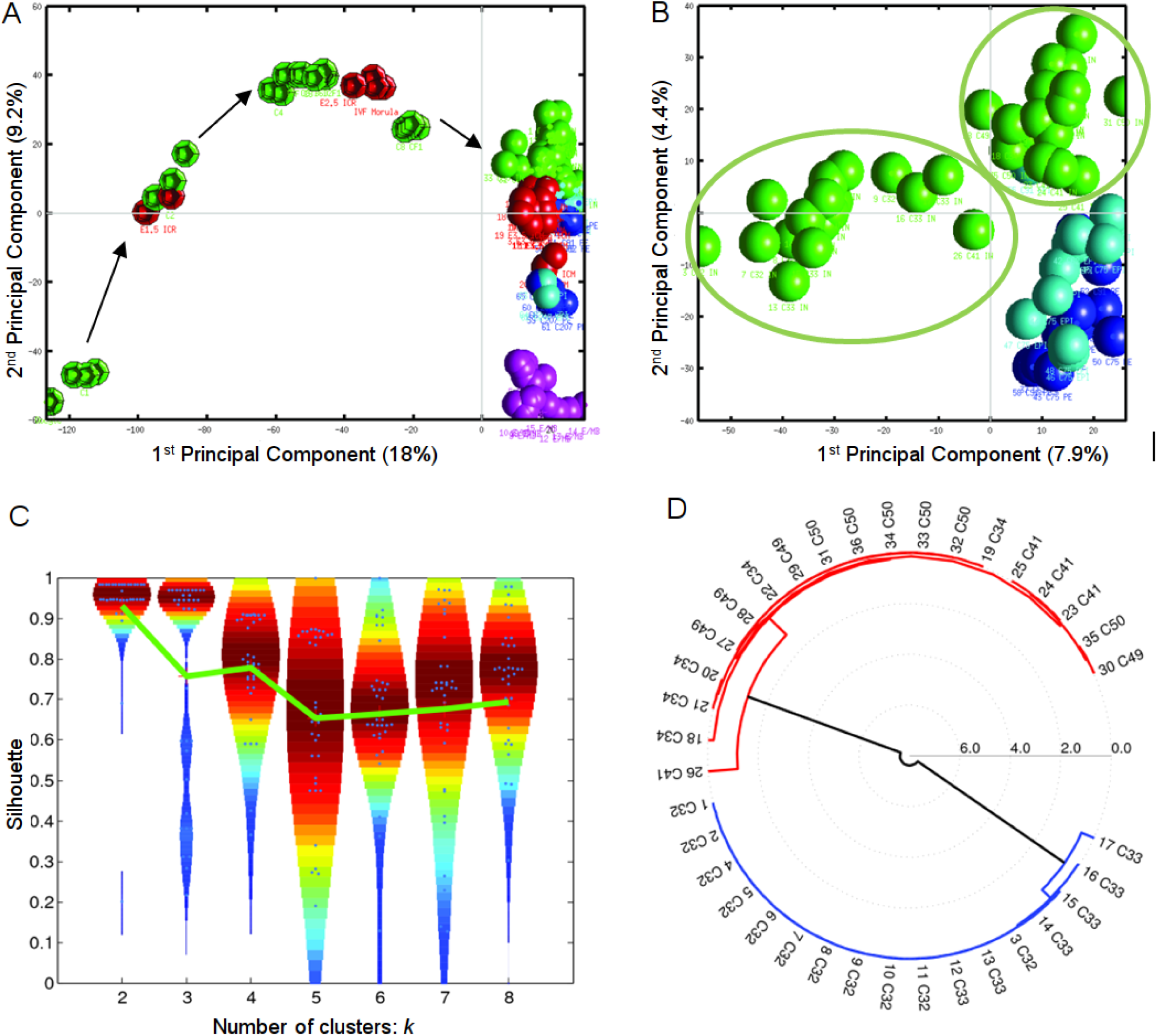
Hierarchical optimal *k*-means clustering (HO*k*M) reveals a splitting into two clusters of the gene expression in the ICM of the mouse embryo at E3.25. Bi-dimensional Principal Component Analysis (PCA) of transcriptomics data from (A) all *in vivo* samples (Table 1) Dodecahedra and spheres mark bulk and single cells, respectively. (B) E3.25 and E3.5 (Ohnishi *et al.*, 2014). The green spheres mark the E3.25 cells, while the light and dark blue spheres mark the E3.5 EPI and E.3.5 PE cells, respectively. The two green ellipses encircle the two groups of E.25 cells, posteriorly classified by our HO*k*M method and named as E3.25-LNCs and E3.25-HNCs. (C) Violin plot of the silhouettes of the HO*k*M trajectories. The green line marks the position of the medium silhouette distributions. (D) Dendrogram of the optimal clustering (*k_o_*=2).

PCA explains only −30% of the transcriptomics information (Fig. S1AB). Therefore, to analyze the splitting event of the E3.25 ICM, accounting simultaneously for maximum amount of information and reducing the intrinsic noise of the single-cell data, we developed a clustering algorithm, termed hierarchical optimal *k*-means (HO*k*M), that determines the optimal number of cell subgroups and their members (see Methods). Ohnishi *et al.* (2014) attempted to find an early splitting event between PE and EPI precursors at E3.25 with a conventional clustering method but detected no such event. With our HO*k*M clustering algorithm, we looked to identify any splitting event, rather than between PE and EPI precursors. The E3.25 ICM cells split best into two groups (optimal silhouette coefficient *k* = 2) (Fig. 1C) based on the number of cells of the embryo of origin: one comprising all cells from LNC (32-33-cell) embryos, and another comprising all cells from HNC (34-50-cell) embryos (Fig. 1D). We call these two clusters E3.25-LNC and E3.25-HNC from here on. Thus, we recognized a possible developmental sub-stage splitting event.

### Cell junction and plasma membrane *Bsg*, *Ctnnb1* and *Fgfr1* genes are highly expressed in E3.25-HNCs

Next, we searched for the specific genes driving the E3.25 ICM optimal clustering performing a Differentially Expressed Genes (DEGs) analysis between E3.25-LNC and E3.25-HNC. We found 399 DEGs for the highly expressed transcripts in E3.25-HNC (HNC-h-DEGs), the top ranked depicted in Fig. 2A and the rest in Fig. S3. The top ranked HNC-h-DEG is basigin *Bsg*/*CDl47*/*EMMPRIN* (Fig. 2A), known to regulate the canonical Wnt/beta-catenin signaling pathway (Sidhu *et al.*, 2010) and thought to be regulated by hypoxia (Ke *et al.*, 2012). The fourth top ranked HNC-h-DEG is the key mediator of the Wnt pathway *Ctnnbl* ({*3-catenin*) (Fig. 2A), which, together with the signal transducer *Ywhaz*, is one of the main hubs of the protein binary interaction network of the HNC-h-DEGs, the major one being Oct4 (Fig. 2B). Among the top HNC-h-DEGs, 15^th^, is *Fgfrl*, required for the lineage establishment and progression within the ICM (Kang *et al.*, 2017).

**Figure 2.**
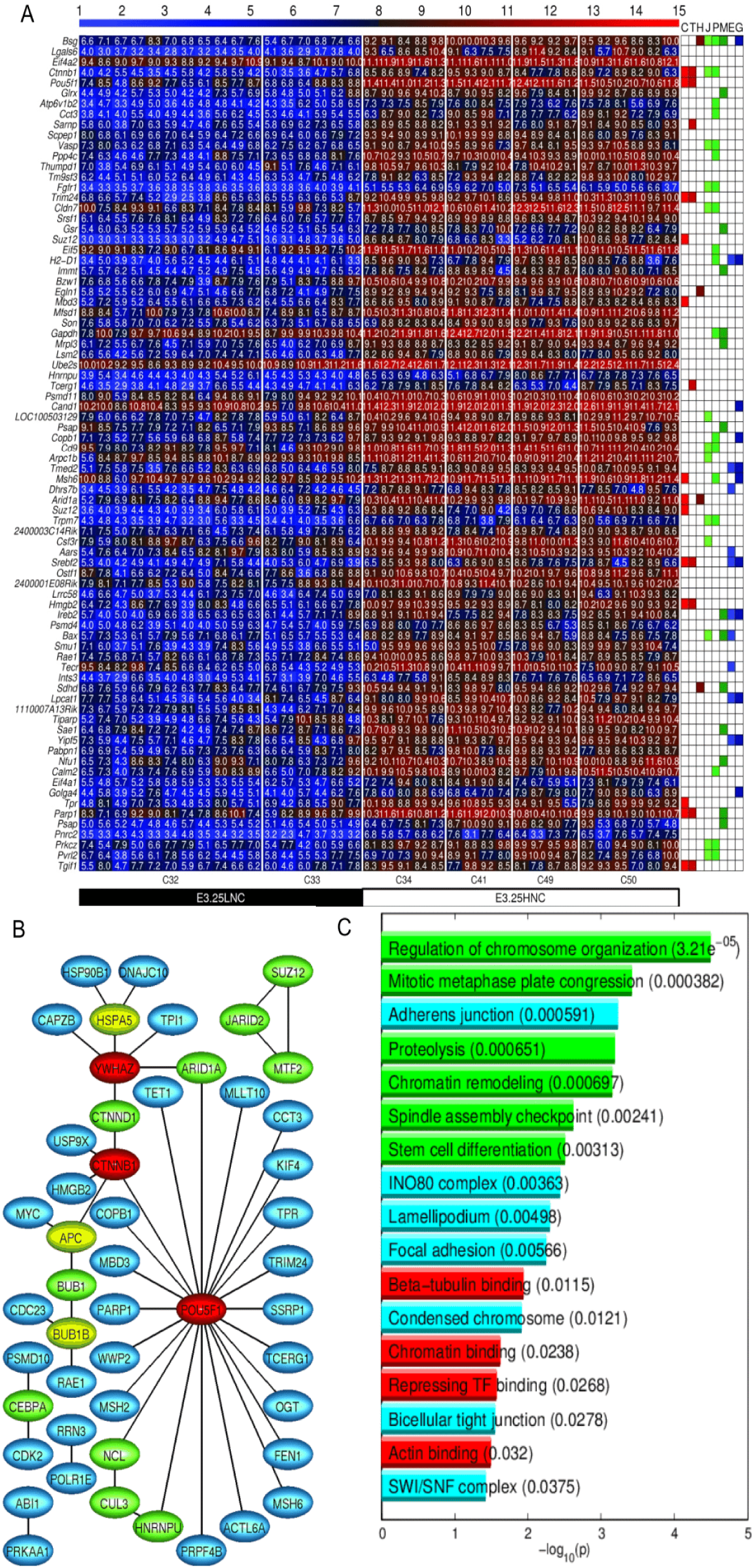
Expression of *Oct4* and several chromatin remodelers is stabilized at high level in E3.25-HNCs. (A) Heatmap of the expression of the 80 top ranked HNC-h-DEGs in decreasing order of significance. The color bar codifies the gene expression in log_2_ scale. Higher gene expression corresponds to redder color. The table to the right annotates GO terms: C (Chromatin remodelers), T (Transcription factor activity), H (Hypoxia), J (Cell junction), P (Plasma membrane), M (Mitochondrion), E (Endoplasmatic reticulum), G (Golgi apparatus). (B) Protein binary interaction network of the HNC-h-DEGs. The node color codifies incidence number (blue, green, yellow and red for incidences 1,2,3 and more than 4, respectively). (C) Bar plot of the -log_10_(*p*-value) of the significant enriched GO terms of HNC-h-DEGs. Longer bars correspond to higher statistical significance of the enrichment (*p*-values inside parentheses). The red, green and cyan bars correspond with molecular function, biological process and cellular compartment GO terms, respectively.

### *Oct4* and several chromatin remodelers expression is stabilized at high level in E3.25-HNC

The Gene Ontology (GO) analysis of the HNC-h-DEGs reveals that among statistically significant enriched GO terms are chromatin remodelers such as the INO80 and the SWI/SNF complex, and cell-cell interaction terms such as adherent junction, focal adhesion and bicellular tight junction (Fig. 2C). A detailed list of all found GO terms (Fig. S5) and their corresponding genes is provided in Tables S1-3. The HNC-h-DEGs involved in chromatin-remodeling complexes together with their roles are enlisted in Table S4 and annotated in Fig. 2A and Fig. S2. Among the top ranked DEGs we discovered one of the pluripotency regulation masters: *Pou5f1* (*Oct4*) that has much lower expression in E3.25-LNCs than in E3.25-HNCs, with some oscillatory expression spikes characteristic of a “salt and pepper” expression pattern. In all E3.25-HNCs, *Oct4* is stabilized at very high expression level. The violin plots of the expression of *Oct*4 across all early developmental stages (Fig. S4) reveal a continuously growing expression since oocyte till 8-cell stage, which is perturbed to lower expression in 32- and 33-cell embryos in order to come back again to high expression during the following developmental stages. The expressions of the *Oct4* common regulators *Sox2* and *Nanog* correlate slightly with that of *Oct4* (Fig. S4). At E3.25 *Nanog* shows the previously reported “salt and pepper” expression pattern, with a small population following the low expression of *Oct4* in the 33- and 34-cell embryos, whereas the expression of *Sox2* remains almost continuously high, with exception of some cells from 32-, 41-, and 50-cell embryos (Fig. S4). The enrichment of pluripotency genes among the HNC-h-DEGs prompted an elucidation of the strength of the pluripotency network in the E3.25-HNCs through an intersection of the HNC-h-DEGs with the interactomes of the master regulator of pluripotency Oct4 and the main pluripotency players (Sox2 and Nanog) (see Methods). Among the proteins shared by the Oct4-Sox2 interactome with the HNC-h-DEGs, we found four genes (Fig. S6): *Cct3, Hnrnpu, Parp1*, and *Ssrp1* whose function is described in Table S5. Particularly, *Hnrnpu* and *Parp1* are *Oct4* transcriptional regulators (Vizlin-Hodzic *et al*., 2011; Roper *et al*., 2014). Among the proteins shared by the Oct4-Nanog interactome with the HNC-h-DEGs we found four common genes (Fig. S6): *Aridla*, *Mbd3*, *Msh6* and *Tet1*, all of them chromatin remodelers (Table S4).

### *Fgf4, Sox2* and *Dvl1* segregate into two different trajectories before E3.25-LNC

We checked for the transitions of the transcriptomics dynamics performing a bifurcation analysis from E3.25 to E4.5 stage (E3.25-LNC, E3.25-HNC, E3.5, E4.5) and determined the first genes associated to dynamic segregation. Such analysis was based on clustering the gene expression of the most highly variable expressed genes across each embryonic day (see Methods). We classified the gene trajectories into 15 categories based on the number of bifurcations of expression for each of the four embryonic stages in order to determine the segregation moment. The four most interesting trajectory categories are: (2-2-2-2) with two bifurcation points at each of the 4 embryonic stages that corresponds to a very early segregation time (before E3.25-LNC), (1-2-2-2) that corresponds to an intermediate segregation time E3.25-HNC, (1-1-2-2) with no bifurcation before E3.5 but bifurcating at E3.5 and remaining so till E4.5, and (1-1-1-2) corresponds to late segregation time E4.5 (Fig. 3).

**Figure 3.**
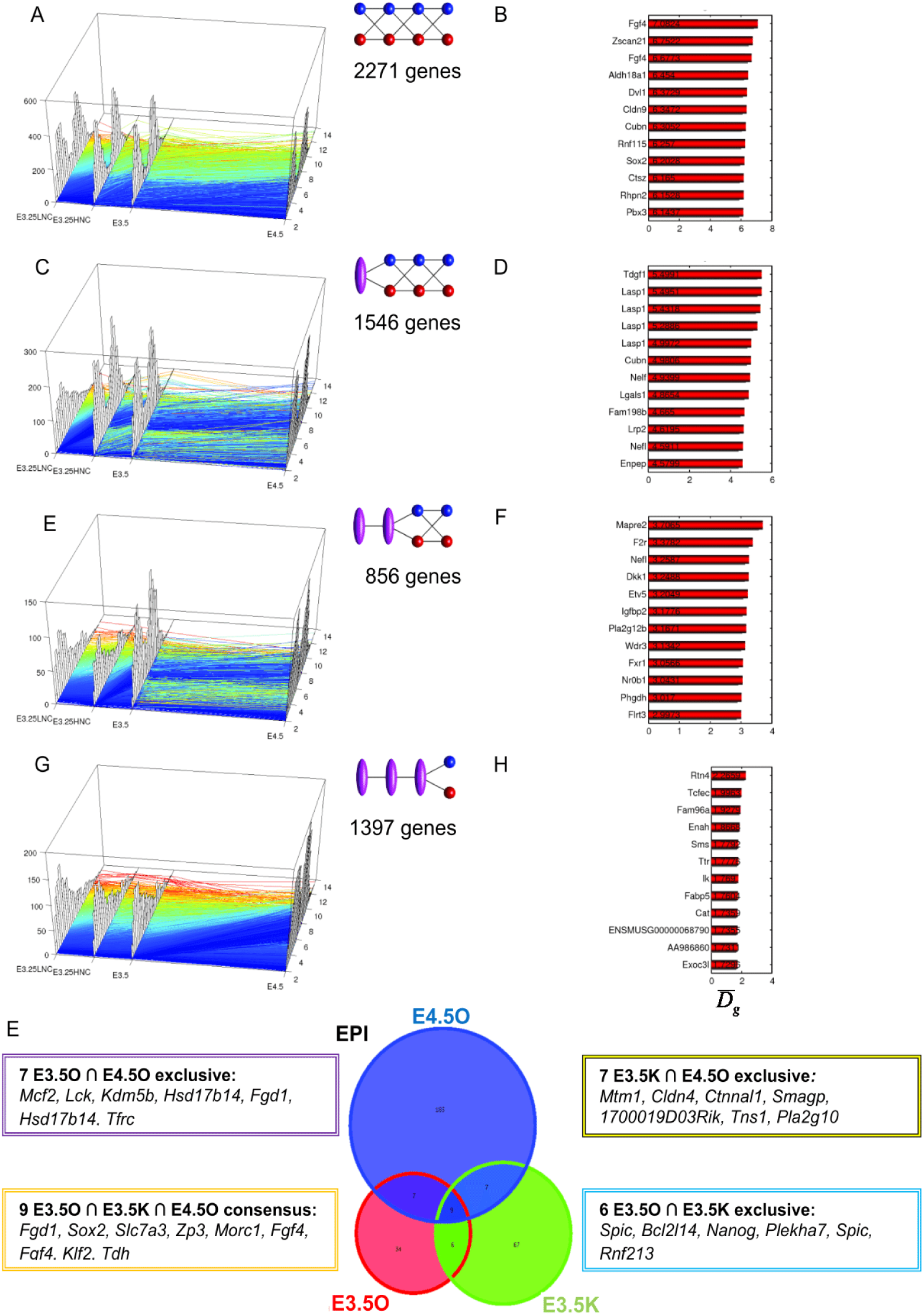
Dynamics of the transcriptomics bifurcations across E3.25-LNC, E3.25-HNC, E3.5 and E4.5. (A), (C), (E), (G) Transcriptomics trajectories ordered from earlier to later bifurcation decision. Each trajectory color corresponds to the level of expression at the earliest time of the trajectory. Bluer color corresponds to lower expression at the earliest measured time. The histogram of the transcriptomics level of the trajectories is shown in the *z*-axis at each embryonic day. The two-point passing stages represent a bifurcation of the gene trajectory at such stage, whereas the one-point passing stage represents a continuous distribution of states that cannot be segregated into two modes. (B), (D), (F), (G) Top 12 transcripts with the maximum separation of the bifurcation of their corresponding trajectories defined by the difference 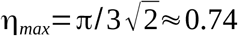 of the bifurcations means.

Interestingly, category (2-2-2-2) with bifurcation time before E3.25-LNC is the most populated with 2271 genes (Fig. 3A). This indicates that the ICM cells that eventually become PE and EPI progenitors at E4.5 carry considerable heterogeneity even before E3.25-LNC. The very top gene with the widest bifurcation from this category is the EPI marker *Fgf4*, with two microarray probes among the 12 top-ranked transcripts (Fig. 3B). The violin plots of the gene expression distribution of *Fgf4* across all the preimplantation developmental stages (Fig. S4) confirm that *Fgf4* shows a splitting pattern ever since the 2-cell stage. Among the top genes in the list is also the *Pou5f1* partner *Sox2*, the member of the Dishevelled family of proteins which regulate both canonical and non-canonical (planar cell polarity pathway) Wnt signaling, as well as *Dvl1* and the PE lineage marker *Cubilin* (*Cubn*).

### E3.25-HNC embryos are more developed than E3.25-LNC

The PCA suggested that E3.25-LNC are more similar to earlier than E3.25 developmental stages, while E3.25-HNC are more similar to later than E3.25 ones (Fig. 1B). To confirm that, we used as representative members of each cluster their top high DEGs between E3.25-LNC *vs* E3.25-HNC, and we analyzed their transcriptomics distributions across other developmental stages using violin plots (Fig. 4A). All the stages before the 33-cell embryos are dissimilar in terms of distribution spread, with more dispersed distributions than after the 34-cell stage. The transcriptomics distributions after the 34-cell stage are similar both in terms of median values and spread (Fig. 4A). The violin plots of the top E3.25 LNC-h-DEGs (Fig. 4B) depict their transcriptomics distributions as unique and not similar to any of the distributions associated to other developmental stages, being slightly closer to the 4-cell stage but surprisingly not to the 8-cell one. Thus, E3.25-HNCs are more related to later developmental stages than E3.25-LNCs.

**Figure 4.**
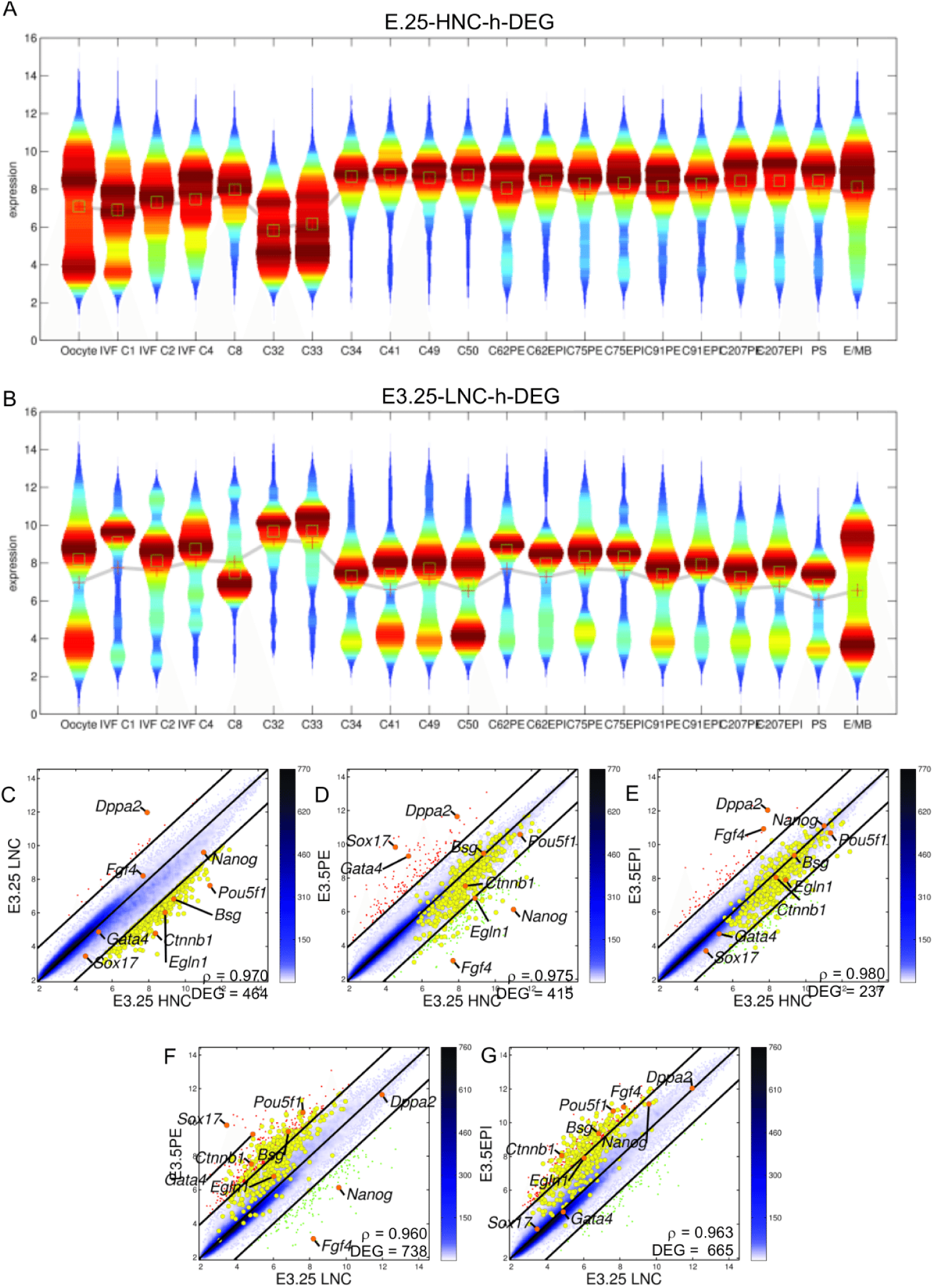
E3.25-HNCs are more developed than E3.25-LNCs. Violin plots of the distribution of the expression of (A) E3.25 HNC-h-DEGs and (B) E3.25 LNC-h-DEGs across different developmental stages. The mean and median are shown as red crosses and green squares, respectively. Pairwise scatter plots of (C) E3.25-LNC *vs* E3.25-HNC, (D) E3.5-PE *vs* E3.25-HNC, (E) E3.5-EPI *vs* E3.25-HNC, (F) E3.5-PE *vs* E3.25-LNC, (G) E3.5-EPI *vs* E3.25-LNC. The black lines are the boundaries of the 2-fold changes in gene expression levels between the paired samples. Transcripts up-regulated in ordinate samples compared with abscissa samples, are shown with red dots; those down-regulated, with green. The positions of some markers are shown as orange dots. The color bar indicates the scattering density. Darker blue color corresponds to higher scattering density. The transcript expression levels are log_2_ scaled. ρ is the Pearson’s correlation coefficient. The E3.25 HNC-h-DEGs are overimposed as yellow dots.

Interestingly, E3.25-HNCs are dissimilar to both PE and EPI (Fig. 4A) based on the top DEGs defining the E3.25 HNCs. Anyway, we checked for a possible global similarity of E3.25-HNCs with EPI or PE using all transcripts and not only the top highly DEGs. We made scatter plots, calculating the average pool of all the cells of E3.25-LNC, E3.25-HNC, E3.5 PE and E3.5 EPI. The split of E3.25-LNC and E3.25-HNC is preserved after performing the average of the respective single cells (Fig. 4C). They have, not very high for transcriptomics studies, correlation coefficient ρ = 0.970 and 464 DEGs. Both E3.25-HNC and E3.25-LNC express in a similar way the *Fgf4* and *Nanog* EPI markers and the *Gata4 and Soxl7* PE markers. E3.25-HNCs are closer in terms of correlation coefficient p and number of DEGs to both PE and EPI E3.5 populations compared to the E3.25-LNCs (Fig. 4FG). It is noteworthy that E3.25-HNCs are close to both PE (ρ = 0.975) and EPI (ρ = 0.980) but not to any of the two lineages specifically. Thus, the higher transcriptomics similarity of the E3.25-HNC embryos with older-than-E3.25 embryos is not related to the cell fate but rather to the developmental stage, early or late.

### E3.25 HNC-h-DEGs share transcriptomics features with E3.5 *Fgf4-*KO

Fgf4 is the main determinant of the EPI-PE segregation (Chazaud *et al*, 2006; Guo *et al*, 2010; Yamanaka *el al.*, 2010). In an attempt to see whether the set of genes defining the HNC cluster, E3.25 HNC-h-DEGs, is related in some way to the PE and EPI cell specification in the ICM, we compared the E3.25 HNC-h-DEGs with *Fgf4* knockouts (*Fgf4*-KO) from Ohnishi *et al*. (2014) (Fig. S7). We found that the HNC-h-DEGs are lowly expressed in E3.25 *Fgf4*-KO and E4.5 *Fgf4*-KO cells, however they are highly expressed in E3.5 *Fgf4*-KO, thus, the transcriptomics profile of E3.5 *Fgf4*-KO resembles the profile of E3.25-HNCs. For the subset of E3.25 HNC-h-DEGs, the E3.25 *Fgf4*-KO cells are similar to the E3.25-LNCs. This suggests that both E3.25 *Fgf4*-KO and E3.25-LNCs have still undetermined ICM fate. The E3.25 *Fgf4-*KOs originate from low and high number-of-cell embryos, which hints that lack of Fgf4 prevents inner cells from even high-number-of-cell embryos (E3.25 *Fgf4*-KO with > 33 cells) to take secure ICM fate and holds back the inner cells at the undetermined ICM cell fate stage (E3.25 with <34 inner cells). Interestingly, the effect of the *Fgf4*-KO at E4.5 reverts the transcriptomics profile based on the subset of the E3.25 HNC-h-DEGs to the E3.25-LNC state, whereas the E3.5 *Fgf4*-KO preserves the E3.25-HNC transcriptomics state.

Additionally, we followed the evolution of the transcriptomics profiles based on the subset of E3.25-HNC-h-DEGs in the PE and EPI cells at E3.5 and E4.5, and compared them with the *Fgf4*-KOs. Three groups of cells (E3.5PE, E3.5EPI and E3.5 *Fgf4*-KO) are very similar in their transcriptomics profile based on the subset of E3.25-HNC-h-DEGs (Fig. S7). The similarity between E3.5 PE and E3.5 EPI is noteworthy and can be related to the fact that at E3.5 the PE and EPI cells can still revert their fate to the opposite fate. The subset of E3.25 HNC-h-DEGs is related to a chromatin remodeling process that pushes forward the more narve pluripotent state (E3.25-LNC) to a more advanced pluripotent state (E3.25-HNC and E3.5 (PE+EPI)) and it seems that *Fgf4*-KO does not affect this process. This is confirmed by the absence of *Fgf4* in the E3.25-HNC-h-DEGs and by the very early bifurcation of *Fgf4* before E3.25. Strikingly, at E4.5 the profile of the E3.25 HNC-h-DEGs in the *Fgf4*-KO cells is similar to the profile of the E3.25-LNC, even though the embryos at E4.5 have very high number of cells, much higher than 33, pointing that Fgf4 affects the machinery that drives E3.25-LNC to more advanced/differentiated pluripotent state in the HNC at E3.25 and E4.5 but not at E3.5.

### E3.25-HNCs are more enriched in PE and EPI markers than E3.25-LNCs

To study whether the E3.25-LNCs and E3.25-HNCs differ in terms of lineage differentiation, we predicted early PE and EPI lineage markers from the consensus of the E3.5 transcriptomics single-cell datasets of Ohnishi *et al.* (2014), E3.5O and E4.5O, and Kurimoto *et al.* (2006), E3.5K. Since the Kurimoto *et al.* (2006) E3.5 ICM dataset was not initially classified into PE and EPI, we applied our HO*k*M algorithm to classify it. We obtained that the optimal number of clusters is *k* = 2, with one cluster comprising cells 3-5, 8, 10, 11, 15, 17 and 20, and another comprising cells 1, 2, 6, 7, 9, 12-14, 18 and 19 (Fig. 5A). For validation purposes, we applied our HO*k*M algorithm to the already classified PE and EPI cells from E3.5 and E4.5 of Ohnishi *et al.* (2014). Even though our HO*k*M algorithm is an unsupervised clustering method, it classified with 100% accuracy the two EPI-PE datasets, namely E3.5O and E4.5O (Fig. 5BC).

**Figure 5.**
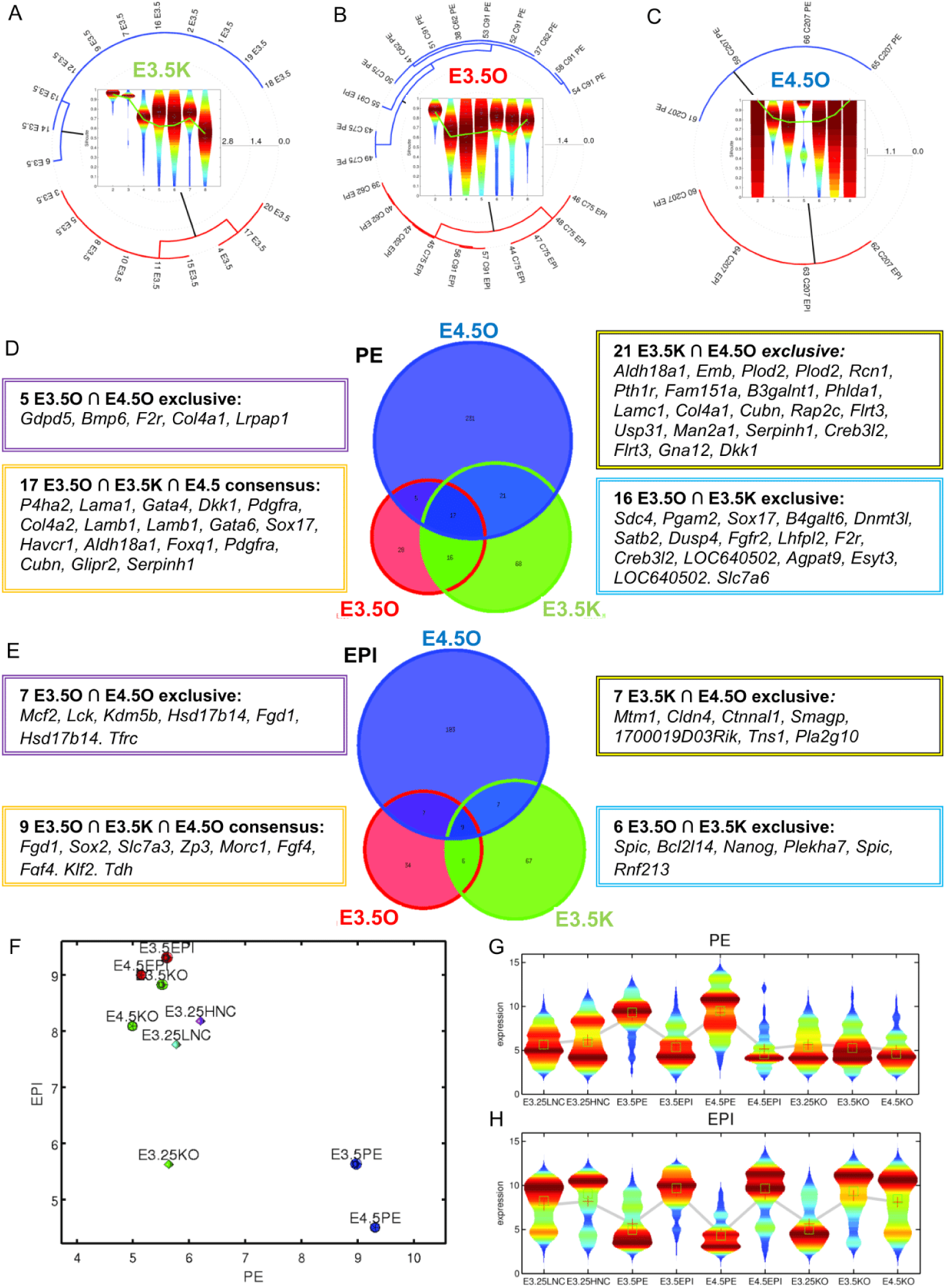
Map of samples on the EPI *vs* PE space built from the consensus of EPI and PE markers predicted through HO*k*M. Polar dendrograms of the HO*k*M of (A) the E3.5 data from Kurimoto *et al.* (2006) E3.5K(urimoto), and from Ohnishi *et al.* (2014) (B) E3.5O(hnishi) (C) E4.5O(hnishi). Violin plots of the silhouettes of the HO*k*M trajectories are presented in the center of each dendrogram. The overimposed green line in the violin plots marks the position of the medium silhouette distributions. Euler-Venn diagrams of the transcripts shared by the (D) PE and (E) EPI populations. (F) Map of single-cell transcriptomics data on the EPI - PE space. Violin plots of the distributions of the consensus (G) EPI and (H) PE markers. The mean and median are shown as red crosses and green squares, respectively.

After the correct EPI-PE classification, we obtained the consensus shared markers. We searched for the statistically significant DEGs between the EPI *vs* PE clusters of each dataset, and performed the intersections of the corresponding lists of estimated PE and EPI markers (Fig. 5DE). Among the genes shared by the three common PE intersections are the known markers *Sox17*, *Gata6* and *Cubn*. Among the E3.5O⋂E3.5K *exclusive* PE intersection corresponding to earlier markers are known markers such as a second probe of *Sox17*, and *Fgfr2*, but also transcripts such as two probes of *LOC640502* that correspond to *Uap1* (UDP-N-acetylglucosamine pyrophosphorylase 1) and that has not been reported as a PE marker. Among the genes shared by the common intersection of the three EPI sets is the known marker *Fgf4* (with two probes). Among the E3.5O⋂E3.5K *exclusive* EPI intersection is the known marker *Nanog*.

The only E3.5-specific PE and EPI markers among the HNC-h-DEGs are *CpneB* and *Ubxn2a*, respectively. The E3.5-specific EPI marker (Gerovska and Arauzo-Bravo, 2015) Ubxn2a (UBX domain protein 2A)/Ubxd4, regulates the cell surface number and stability of alpha3-containing nicotinic acetylcholine receptors (Rezvani *et al*, 2009) and is located in both the endoplasmic reticulum and cis-Golgi compartments interacting with the ubiquitin-proteasome system. The calcium-dependent membrane-binding phospholipid protein Cpne8 may regulate molecular events at the interface of the cell membrane playing a role in membrane trafficking.

We built a 2D map of the EPI-PE space based on consensus predicted markers, assigning to each cell the transcriptomics mean across all PE and EPI markers (Fig. 5F). All wild type PEs, except one, locate as high PE-low EPI. E3.25-LNCs and E3.25-HNCs are mainly high EPI - low PE, where E3.25-HNCs are with slightly higher PE and EPI expression. In the *Fgf4*-KO case, E3.25 KOs are depleted in both PE and EPI markers (low PE-low EPI), while E3.5 KOs and E4.5 KOs are low PE-high EPI. The violin plots showing the variability of the expression of the PE and EPI markers for each cell type confirm that the E3.25-HNC express slightly higher both PE and EPI markers than the E3.25-LNC, while the *Fgf4*-KOs express scarcely PE markers and highly EPI markers at E3.5 and at E4.5 but not at the early E3.25 stage (Fig. 5GH).

### E3.25-HNC transcriptomics profiles are closest to extreme na1ve ESC compared to all other ESCs

To assess the pluripotency fingerprint of the E3.25-E4.5 inner cells of Ohnishi *et al.* (2014), we compared their transcriptomics profiles with those of all the available ESCs in the GEO database from Affymetrix Mouse Genome 430 2.0 Arrays. The *in vitro* ESCs cluster away from the *in vivo* E3.25-E4.5 inner cells (Fig. 6A). To find the closest *in vitro* pluripotent counterparts of each developmental stage from E3.25 to E4.5, we averaged the single-cell transcriptomics profiles of each stage and calculated the distance to each ESC profile. The shortest distance between the set of E3.25-E4.5 inner cells and the ESCs is between the E3.25-HNCs and the extreme narve 2i+LIF ESCs (Fig. 6B). The developmental transition from 33 to 34 cells (E3.25-LNC to E3.25-HNC) decreases dramatically the distance with the extreme narve ESCs. Most of the HNC-h-DEGs are similarly expressed in the E3.25-HNCs and the 2i+LIF ESCs (Fig. 6C). The egl-9 family hypoxia-inducible factor 1, *Egln1*, is among the few HNC-h-DEGs down-regulated in the 2i+LIF ESCs (Fig. 6C).

**Figure 6.**
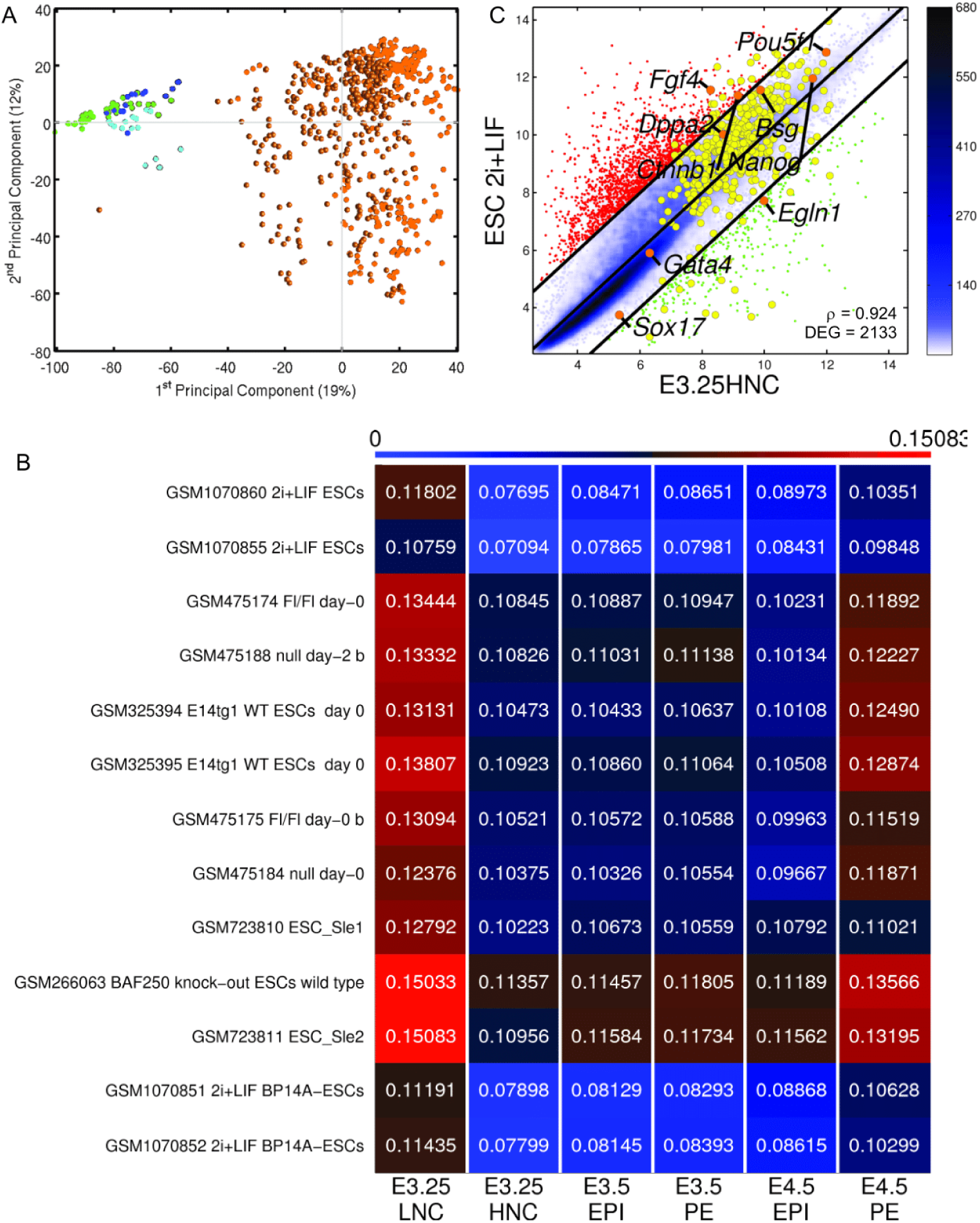
Comparison of E3.25-E4.5 stage inner cells (Ohnishi *et a/.*, 2014) with all existing ESC Affymetrix Mouse Genome 430 2.0 Array transcriptomics profiles. (A) PCA. Green circles and dodecahedra mark E3.25 LNCs and HNCs, respectively. Cyan circles and dodecahedra mark E3.5 and E4.5 PE, respectively. Blue circles and dodecahedra mark E3.5 and E4.5 EPI, respectively. Orange circles mark ESCs. (B) Heatmap of the Spearman correlation distances of the top closest ESC to the different preimplantation developmental stages. (C) Pairwise scatter plots of ESC 2i+LIF *vs* E3.25-HNC. The black lines are the boundaries of the 2-fold changes in gene expression levels between the paired samples. Transcripts up-regulated in ordinate samples compared with abscissa samples, are shown with red dots; those down-regulated, with green. The positions of some markers are shown as orange dots. The color bar indicates the scattering density. Darker blue color corresponds to higher scattering density. The expression levels are log_2_ scaled. p is the Pearson’s correlation coefficient. The E3.25 HNC-h-DEGs are overimposed as yellow dots.

### ScRNA-seq study shows that *Oct4* expression is stabilized at high level in the ICM of late stage 32-cell embryos

To validate our findings based on microarrays, we searched for a single cell RNA-seq dataset that encompasses the developmental time around E3.25 and found that of Posfai *et al.* (2017), GSE84892. Unfortunately, the information on the exact number of cells of the embryos from the early and late 32 cell stages, from which the single cells for the RNA-seq analysis were dissected is not preserved (personal communication). However, (1) The early and late 32-cells have been harvested 72h and 78h post fertilization, and correspond to E3.0 and E3.25, respectively, (2) the early and late 32 cell stage differ in the degree of formation of blastocoel, and (3) Most of the early 32 TE cells are “specified and committed and few cells are capable of forming ICM” and among the late 32 cells, “all TE cells are specified and committed", and for both early and late 32 cells, “ICM specified, but not committed, TE fate can be induced by forced outside exposure or by inactivating Hippo signalling” (Figure 7 in Posfai *et al.* (2017)). This situation resembles what we observed from our analysis of the data set of Ohnishi *et al*. (2014) that “E3.25-HNC embryos are more developed than E3.25-LNC” but not yet committed to PE or EPI lineage. Therefore, we searched for the DEGs between the early and late 32 cell stage ICM cells from Posfai *et al.* (2017). The top 80 upregulated genes in the late 32-cell ICM (L32ICM-h-DEGs) are shown in the heatmap of Fig. 8, together with their expression in the ICM cells during early and late 16 cell stage and in the ICM of the 64 cell stage of Posfai *et al.* (2017). The intersection between the top 80 L32ICM-h-DEGs and the top 80 HNC-h-DEGs, includes *Pou5f1/Oct4*, *Eif4a2* and *Trim24*, all of them among the several top ones in both L32ICM-h-DEGs and the HNC-h-DEGs. The expression of *Oct4* in the ICM RNA-seq data from Posfai *et al.* (2017) follows that of the Affymetrix array data presented in the *Oct4* violin plot in Supplementary Fig. S4. *Oct4* is higher expressed and stabilizes at high level in the late 32 ICM cells (Fig. 2A and Fig. 8). Before the 32 cell stage, *Oct4* is high and stable in both microarray and RNA-seq data (Supplementary Fig. S4 and Fig.7). Interestingly, together with *Oct4*, among the top 80 L32ICM-h-DEGs, we found the core pluripotency factors *Klf4* and *Sox2*, as well as the pluripotency associated transcript 9, *Platr9/*2410114N07Rik. Additionally, we found *Gata6*, observed in the majority of ICM cells in early blastocysts and later confined to the PE progenitors at the mid-blastocyst stage (Chazaud *et al.*, 2006). We have checked the expression of L32ICM-h-DEGs in the ENCODE project and found that these genes have the highest expression in either testis adult (*Trim24*, *Pcbp2*, *Aven*, *1B10044D09Rik*, *Csd/Ybx3*, *Pou5f1*, *Yif1b*) and ovary adult (*Yy1*, *Pum1*), or placenta adult (*Morf412*,*Trap1a* with expression restricted to placenta, *Snx5*), i. e. are genes expressed in either the germ cells of the embryo proper or the extraembryonic placenta. Analogously to the HNC-h-DEGs, we performed an intersection of the L32ICM-h-DEGs (fold-change 1.5) with the interactomes shared by the Oct4-Sox2 and by Oct4-Nanog (Figure S8). Among the proteins shared by the Oct4-Sox2 interactome with the L32ICM-h-DEGs, we found two genes (Fig. S8): *Parp1*, and *Ssrp1*, both of them common with the intersection of the HNC-h-DEGs with the Oct4-Sox2 interactome. Among the proteins shared by the Oct4-Nanog interactome with the L32ICM-h-DEGs we found five common genes (Fig. S8): *Cnot1*, *Esrrb*, *Mbd3*, *Sall4* and *Sox2*, with *Mbd3* being common with intersection of the HNC-h-DEGs with the Oct4-Nanog interactome. *Cnot1*, CCR4-NOT transcription complex, subunit 1, is a component of the core pluripotency circuitry conserved in mouse and human ESCs (Zheng *et al.*, 2012). *Sall4*, spalt like transcription factor 4, is an important pluripotency factor that is required for stem cell maintenance. *Essrb*, estrogen related receptor, beta, enhances induction of naive pluripotency, when transduced with *Oct4*, *Sox2*, and *Klf4* (OSK) into murine fibroblasts (Sone *et al.*, 2017). *Esrrb* is shown to unlock silenced enhancers for reprogramming to naive pluripotency (Adachi *et al.*, 2018).

**Figure 7.**
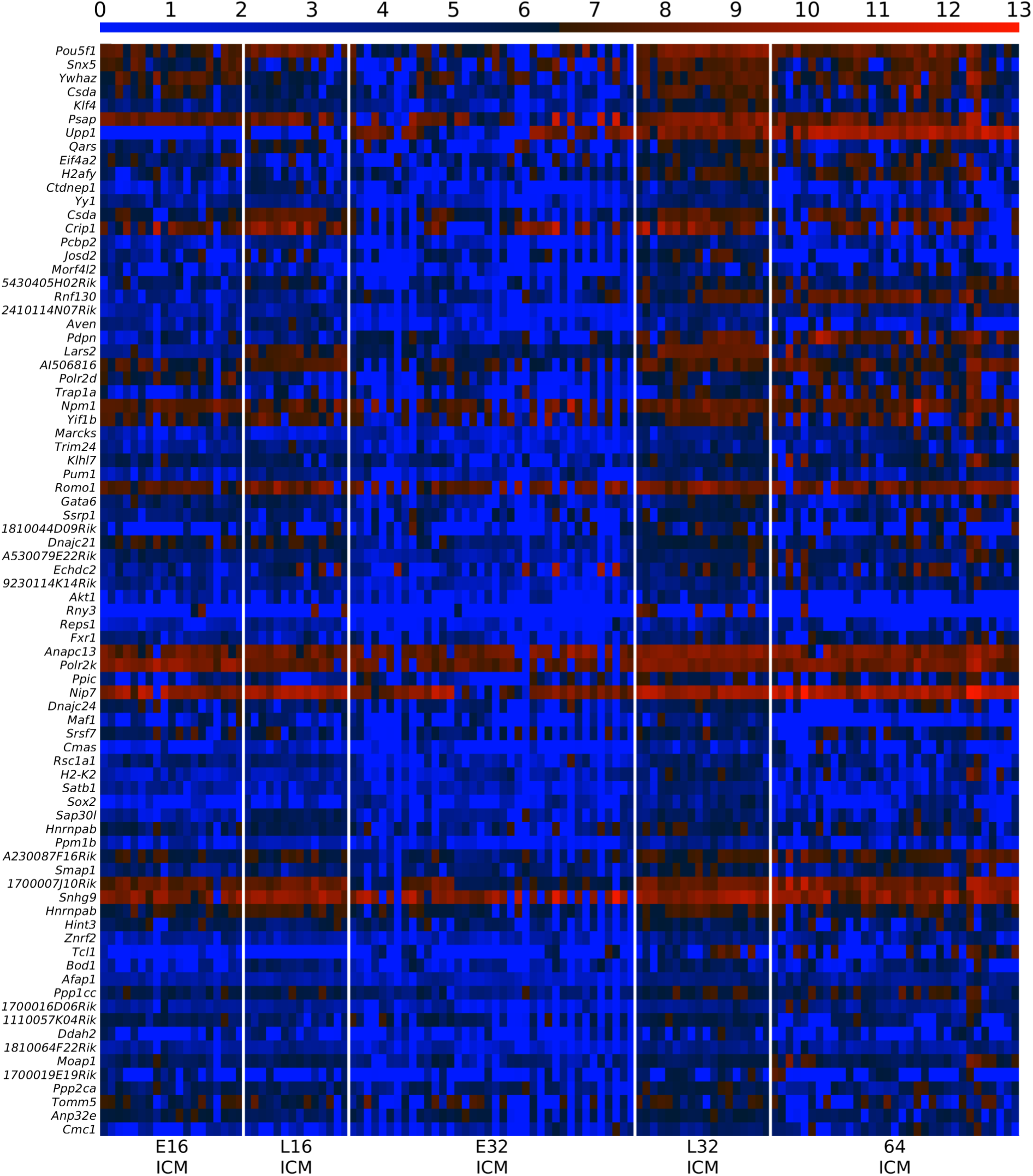
Expression of *Oct4* is stabilized at high level in the late 32-cell ICM. Heatmap of the expression of the 80 top ranked L32ICM-h-DEGs in the ICM cells from early 16 cells to 64 cells from the dataset of Posfai *et al.* (2017). The color bar codifies the gene expression in log_2_ scale. Higher gene expression corresponds to redder color.

**Figure 8.**
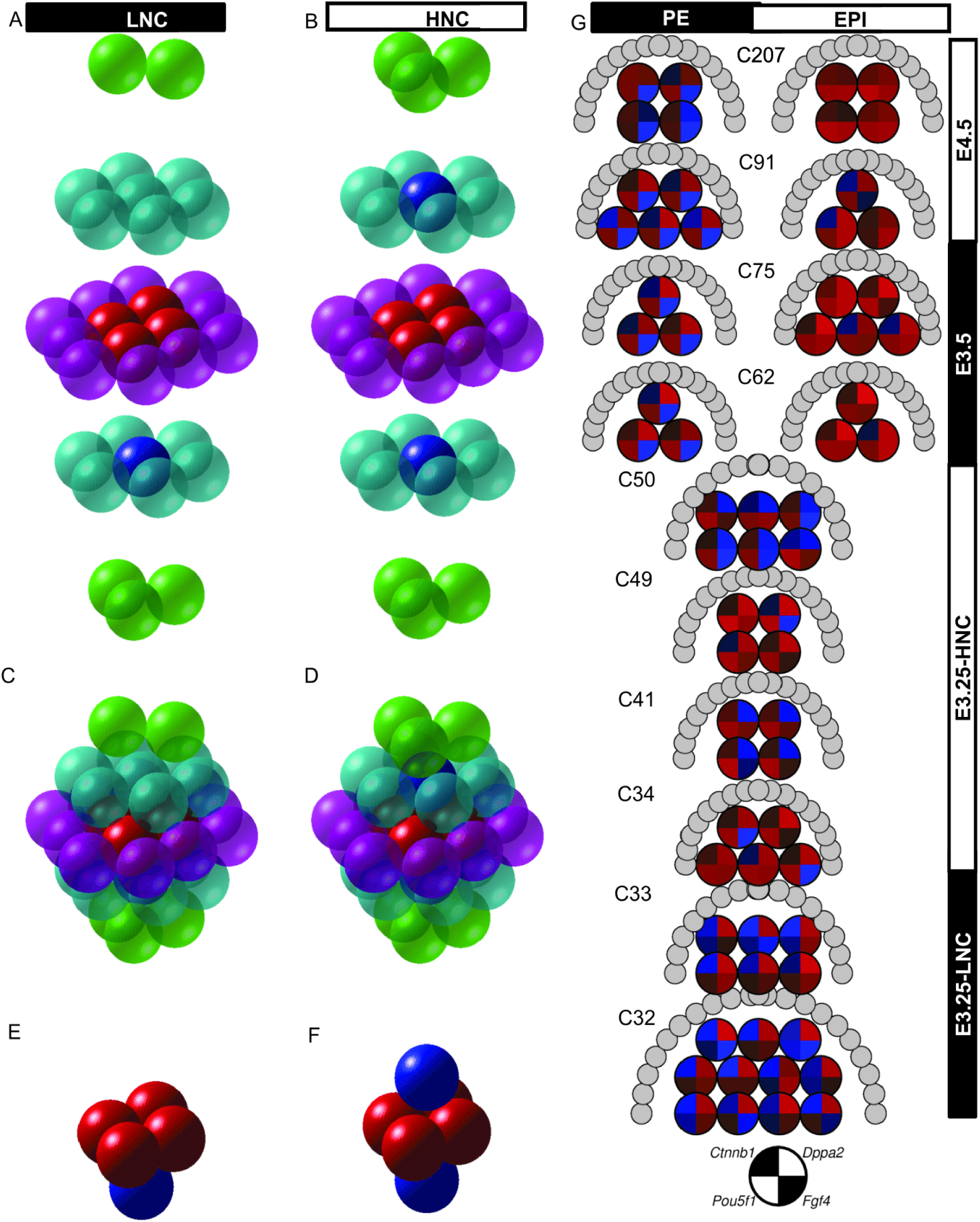
Simplified spatio-temporal mouse embryo model. 3D embryo model based on hexagonal close-packing for embryos with 33 and 34 cells. Layer reconstruction of (A) 33-cell and (B) 34-cell embryos. Packing of the (C) 33-cell and (D) 34-cell embryos. (E) 5-cell and (F) 6-cell kernels formed of cells without external contacts. The transparent and solid spheres represent external and kernel cells, respectively. (G) Temporal representation of ICM cells from blastocysts at different number-of-cell stages and for different cell types. The gene expression of each cell is represented by a pie-chart of the expression of *Pou5f1*, *Fgf4*, *Ctnnbl* and *Dppa2*. The gene expression is represented in log_2_ scale. Higher gene expression corresponds to redder color, and lower expression to bluer color. The genes and their corresponding positions in the pie-chart are represented by the black and white pie-chart in the bottom. The semicircle of gray circles surrounding the top part of each group of single cells represents the blastocyst outer cells.

### Minimum 34 cells are required for a kernel without contacts with the outer shell

We found a clear cut boundary in the transcriptomics profile of the ICM cells from LNC and HNC blastocysts at E3.25. Several of the top 80 HNC-h-DEGs are cell-cell signaling (*Cnntb1*, *Cldn7, Srsf1*), plasma membrane proteins (*Bsg, Cd9, Tmed2*), or/and hypoxia-inducible (*Bsg*, *Sdhd*, *Egln1*) (Fig. 2A). A comprehensive annotation of all HNC-h-DEGs is given in Fig. S3. Therefore, we expect that the strong transcriptomics change arising in embryos with 34 cells are due to the cell signaling emerging from the cell packaging. After compaction at the 8-cell stage the blastomeres are not spherical anymore but resemble a rhombic dodecahedron, however to study how the blastomeres pack, we approximate them by spheres. We built compact models under the hypothesis that the embryo at E3.25 is approximately a sphere with all its cells packed compactly to minimize the embryo volume. Using a random close packing (rcp) the expected packing density is η≈0.64 (Torquato *et al.*, 2000), whereas the demonstration of the Kepler conjecture (Hales, 2005) shows that the optimal arrangement of any packaging of spheres has a maximum average packing density 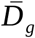. Such density is achieved using hexagonal close-packed (hcp) or face-centered cubic (fcc) regular lattices. The maximum number of near neighbors (coordination number) of hcp and fcc regular lattices is 12. Using a hcp model to fill a spherical embryo, we found that for blastocysts with ≤33 cells, the kernel of cells without contacts with external cells consists of a maximum of 5 cells, while for all blastocysts with ≥34 cells, the kernel of cells without contacts with external cells has a minimum of 6 cells (Fig. 8). The 5-cell kernel forms a pyramid with the 4 base cells having a coordination number (within the other kernel cells) of 3, and the vertex cell having a coordination number of 4; thus, there are 4 x 3 + 4 = 16 contacts (within kernel cells), that, since the maximum coordination number is 12, implies that 16/(5×12) = 26.7% of the kernel contacts are with kernel cells, and 73.3% with inner but not kernel cells. The 6-cell kernel forms an octahedron whose cells have a coordination number (within the kernel cells) of 4; thus, there are 4 x 6 = 24 contacts, which implies that 24/(6×12) = 33.3% of the kernel contacts are with kernel cells, and 66.7% with inner but not kernel cells. These results suggest that to stabilize *Oct4* expression at high level and to activate the epigenetic (DNA unmethylation and chromatin remodeling events), it is necessary to have a kernel of cells with at least a 33.3% of the contacts established inside the kernel; thus, suggesting a minimal interaction competitive cell network in which a third of the internal contacts counteracts the signals from non-kernel cells. Therefore, we hypothesize that in such networks the signal among the kernel cells should be at least twice more intensive than that coming from the outside for the kernel cells. Ctnnb1, Pou5f1 and Dppa2 are the driving forces in the embryo development at E3.25 and in the segregation of the E3.25 ICM cells into groups corresponding to two different developmental stages (Fig. 8G).

## Discussion

Our new hierarchical optimal *k*-means algorithm enabled us to identify two groups of ICM cells during the 32-64-cell embryonic stage corresponding to two subsequent developmental stages with different degrees of potency, rather than to the lineage progenitors PE or EPI. The genes defining these stages indicate that during the cell divisions happening between the 32- and 64-cells, the development of the embryo to 34 cells triggers a dramatic event in which *Oct4* expression is finally stabilized at high level in the ICM to establish the pluripotent state and initialize the chromatin remodeling program. Here we propose that the stabilization of *Oct4* to high expression is a non-cell autonomous process that requires an “inside” environment, i.e. a minimal number of four inner cell contacts in conjunction with a heterogeneous environment formed by the niche of a kernel of at least six inner cells covered by a crust of outer TE cells. The involvement of such non-autonomous cell process prevented the uncovering of the mechanism of establishing of early embryonic pluripotency until now.

We hypothesize that at 34-64 cells (E3.25-HNC) compaction and adhesion events are intensified compared to 32-33 cells (E3.25-LNC) since the HNC-h-DEGs are enriched in glycoproteins such as *Bsg, Psap*, *Cd9*, *Epcam*, *Scarb2, Sep15* and *Dnajc10* and as it has been reported, cell surface glycoproteins play an important role in the development of preimplantation mouse embryo during compaction and trophoblast adhesion (Surani *et al.*, 1981). In the same line, several E3.25 HNC-h-DEGs such as *Ctnnb1*, *Ccni* and *Epcam* are coding cell-adhesion related proteins (Kim *et al.*, 2014; Hagmann *et al.*, 2015), and some are coding transmembrane proteins such as *C1dn7*, *Tm9sf3*, and *Mbtps1*. Additionally, the creating of the “inside” environment at the 34-64-cell stage activates hypoxia signaling, shown by the up-regulation of several hypoxia-inducible genes (*Bsg*, *Acaa2*, *Sdhd*, *Atp1b1*, *Egln1*, *Hsp90b1*).

When the embryo reaches 34 cells, the cell-cell contacts between kernel cells increase to 33.3%. This is also the stage when *Ctnnb1* is highly expressed, indicating activation of the Wnt pathway. It is known that Oct4 is downstream of the Wnt pathway, being a direct target of Ctnnb1 (Li *et al.*, 2012). Wnt signaling promotes maintenance of pluripotency by leading to the stabilization of Oct4 (Kelly *et al.*, 2011), and through interaction with Klf4, Oct4 and Sox2, Ctnnb1 enhances expression of pluripotency circuitry genes promoting cellular reprogramming (Zhang *et al.*, 2014). Wnt/β-catenin triggers the activation of *Oct4* expression since it enhances iPSCs induction at the early stage of reprogramming, however, Wnt/β-catenin is not required for pluripotent stem cell self-renewal (Zhang *et al.*, 2014). We observed that *Oct4* expression is stabilized in E3.25-HNCs, together with activation of several chromatin remodelers. Additionally, a negative regulator of Wnt is Dkk1 (Dickkopf1), which we found among the 22 common PE markers for E3.5 and E4.5 and which is a feedback target gene for Wnt. Among the genes whose expression follows the activation of the Wnt pathway and reported in the literature (www.stanford.edu/-rnusse/wntwindow.html) are also *Sox17* (in the intersection of E3.5 and E4.5 PE markers), *Fgf4* and *Nanog* (in the intersection of E3.5 and E4.5 EPI markers), as well as *Son* which is a E3.25 HNC-h-DEG.

The E3.25-LNC have “salt and pepper” expression pattern of *Oct4*. They have still undetermined ICM fate whereas the E3.25-HNC have robust expression of *Oct4* and confidently taken ICM fate. Such observation has been confirmed through the reanalysis of the scRNA-seq data of early and late 32-cell ICM. The E3.25-HNCs are similar to both E3.5 PE and E3.5 EPI, which is consistent with the fact that E3.25-HNCs are ICM and therefore pluripotent, however expressing simultaneously PE and EPI markers. The late 32-cell ICM cell have upregulated genes high-expressed in both adult testis and ovary, and placenta, thus they are have determined neither EPI nor PE fate. The E3.25-HNCs are also ICM cells that still have not taken the EPI or PE fate. *Dppa2*, highly expressed E3.25-LNC is an early marker of reprogramming to iPSCs involved in the early stochastic reprogramming phase (Buganim *et al.*, 2012). In E3.25-LNC, *Dppa2* it is highly expressed. The dichotomy between high expression of *Dppa2* in the E3.25-LNC and “salt and pepper” expression pattern in E3.25-HNC, and the converse expression pattern of *Oct4*, could produce different binding patterns in preimplantation embryos at E3.25 as it has been observed in ESCs, where Dppa2 binds promoters but is absent from enhancers (Engelen *et al.*, 2015) binding predominantly to promoters with low activity (high enriched in H3K4me3 and low enriched in H3K27), with a promoter binding pattern not correlated with H3K27ac, featured only shared with Kdm5b and Polycomb repressor proteins. These features are different from other pluripotency-inducing factors, such as Oct4 and Esrrb, which predominantly bind moderately active enhancers.

Our comprehensive scanning of all the ESC Affymetrix Mouse Genome 430 2.0 Array transcriptomics profiles in GEO database revealed that the closest *in vitro* counterpart of the *in vivo* E3.25-HNCs are the 2i+LIF ESCs, which constitute the extreme narve end of the pluripotency spectrum existing in the GEO database, not excluding the discovery of closer *in vitro* counterparts in the future.

Concluding, we propose a model to explain the establishment of pluripotency from totipotency at the preimplantation based on a non-cell autonomous process in which when the embryo reaches 34 cells at E3.25, the 33.3% of the cell-cell kernel contacts reach a critical surface to activate the canonical Wnt pathway, which stabilizes its downstream Oct4 target, that in its turn activates the chromatin remodeling to promote the expression of the pluripotency circuitry genes. The transcriptomics heterogeneity underlying the PE-EPI segregation is running well before the 32-cell stage (E3.25-LNC). Both PE (*Cubn*) and EPI (*Fgf4*, *Sox2*) markers, among 2271 genes in total, follow bifurcated trajectories ever since before E3.25-LNC. We propose that *Oct4* stabilization combined with the broad, already existing heterogeneity leads to the distinguishable PE and EPI transcriptomics profiles of the two lineages at E3.5.

## Methods

### Microarray data processing

We collected data from the Affymetrix Mouse Genome 430 2.0 Array platform from bulk and single cell analysis from the GEO and ArrayExpress databases. The annotation records of the collected data are presented in Table 1. The data were normalized with the Robust Multi-array Analysis (RMA) (Irizarry *et al.*, 2013) using the BioConductor software package (www.bioconductor.org). The PCA and the hierarchical clustering of genes and samples was performed with one minus correlation metric and the unweighted average distance (UPGMA) (also known as group average) linkage method.

### Hierarchical optimal *k*-means (HO*k*M) clustering

To account simultaneously for maximal amount of information and reduction of the intrinsic noise of the single-cell data, we developed a clustering algorithm that determines in an optimal way the number of clusters of cells and the cells belonging to each cluster. The algorithm is based on the generation for each tentative number of clusters *k* of a collections of subgroups of samples for different thresholds of noise filters and different number of replicates of the classical *k-*means clustering algorithm that generates a “fingerprint” for each number of clusters *k.* Such “fingerprints” are clustered through a hierarchical clustering algorithm, and the fitness of each cluster is evaluated through the silhouette coefficient.

To eliminate low responsive signal, for each group of single-cell transcriptomics datasets we filtered out all the probes whose maximal signal across all the samples was bellow a detection threshold θ*_d_* = 5. To eliminate low variable signals, for a response threshold θ*_r_*, 2 ≤ θ*_r_* ≤ 8, we filtered out all the probes whose difference between maximal and minimal signal across all the samples was more than S*_r_*. Thus we progressively created sets of probes with higher signal to noise ratio. For each filtered set of probes we took a collection of tentative number of clusters *k*, 2 ≤ *k* ≤ 9, and we applied the *k*-means clustering algorithm for each *k* using 200 different random starting means of clusters, which produced a *k*-partition of the data. Thus, for each *k* we created a collection of partitions (with different S*_r_* and different starting means of clusters). Such collection of partitions is a “fingerprint” of *k.* Next we performed over each “fingerprint” a hierarchical clustering with a standardized Euclidean metric and a Ward (inner squared distance, minimum variance algorithm) linkage method. The dendrograms produced by the hierarchical clustering were cut at the distance that produces *k* clusters. The datasets belonging to each of the *k* clusters were taken as the *k* partition of the data. For such partition we calculated the silhouette coefficient *s_i_* for each dataset *i*, which is a measure of how similar that dataset is to datasets in its own cluster compared to datasets in other clusters, −1 ≤ s*i* ≤1:

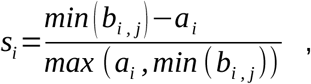

where *a_i_* is the average distance from the *i-*th dataset to the other datasets in its cluster, and *b_i,j_* is the average distance from the *i-*th dataset to datasets *j* in another cluster (Rousseeuw 1987). The silhouette coefficient *S_k_* for the *k* partition is the average of the silhouette coefficient *s_i_* of all datasets. We took as an optimal number of clusters *k_o_*, the *k* that produces the highest *S_k_*, and as optimal clustering, the partition produced by the hierarchical clustering of *k*_o_.

We validated the HO*k*M algorithm by applying it to the already classified PE and EPI cells from E3.5 and E4.5 of Ohnishi *et al.* (2014) and classified the cells with 100% accuracy.

### Transcriptomics dynamics through bifurcation analysis

To eliminate low responsive signals, for each group of single-cell transcriptomics datasets we filtered out all the probes whose maximal signal across all the samples was below a detection threshold θ_*d*_ = 5. To eliminate low variable signals, we filtered out all the probes whose difference between maximal and minimal signal across all the samples was over the response threshold θ_r_ = 8. Thus, we selected a set of 16647 responding probes *RP* with higher signal to noise ratio and reduced the number of probes to be processed in the subsequent stages. For each of the selected probes, and for each group of datasets of the embryonic stages *ES* = {E3.25-LNC, E3.25-HNC, E3.5, E4.5}, we performed a hierarchical clustering with the Euclidean metric and the single (shortest distance) linkage method. For each of the resulting |*RP*|×|*ES*| = 16647×4 = 66588 hierarchical clustering the clustered tree was cut into not more than *k_max_* branches. Since we were interested here in the analysis of two-fate decisions we performed a bifurcation analysis, thus, we used *k_max_* = 2. Each probe *p* was represented by the mean of the data of each cluster. To avoid spurious bifurcations, when the mean values of two clusters were closer than a proximity threshold θ*_d_* = 3, the two clusters were fused into a unique one, and the cluster mean was recalculated as the mean of all the samples. This process was repeated for the four *ES* datasets. Thus, we classified each probe for each embryonic stage *ES* into one or two clusters which were represented by the average of probes belonging to them. The representative value of each probe in each cluster was set by the average of the expression values of the probe in all the cells belonging to that cluster. Since we have *d* = |*ES*| = 4 passing times, *d_i_*⊂*ES* = {E3.25-LNC, E3.25-HNC, E3.5, E4.5}, and one to two bifurcations of transcriptomics values for each passing time, the total number of possible transcriptomics trajectories is (*k*_max_)^*d*^ – 1=15. For each probe *p* belonging to these 15 trajectory categories, and for each passing time *d_i_*, we calculated the difference *D_p_* of the bifurcations means. In case of no bifurcation (*k* = 1) at *d_i_, D_p_* was assigned a zero. The averaged difference 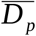 across the *d* passing times was used to rank the transcripts in decreasing order of bifurcation for each of the 15 trajectory types.

### Selection of statistically significant DEGs

To find the statistically significant DEGs between two groups of samples, we calculated the mean of each probe across all the samples of the group. Next, we filtered out all the probes whose absolute value of difference of mean values between the two groups was less than a selection threshold S*_DEG_* = 4 (that corresponds to a fold change of 2 in log_2_ scale), and we applied the Student’s *t*-test. The multitest effect influence was tackled through control of the False Discovery Rate (FDR) using the Benjamini-Hochberg method for correcting the initial *p*-values with significance threshold a_*DEG*_ = 0.001.

### GO statistical significant enrichment analysis of DEGs

The GO terms were taken from the curated collection of molecular signatures (gene set collection C5) of version 3.0 of the Molecular Signatures Database (MSigDB) (Subramanian *et al.*, 2005). The significance of DEGs was analyzed using an enrichment approach based on the hypergeometric distribution. The significance (*p*-value) of the gene set enrichment was calculated using the hypergeometric distribution with significance threshold a_*GO*_ = 0.05.

### Building of the interaction network of the E3.25 HNC-h-DEGs

For the list of E3.25 HNC-h-DEGs we took from the BioGRID database (Stark *et al.*, 2006) (3.4.125 release) all the known binary interactions involving simultaneously two members of the E3.25 HNC-h-DEGs. Then we used the neato layout program of the GraphViz graph visualization software (http://www.graphviz.org/) to calculate an approximation of the interaction network layout. The network was streamlined using in-house implemented functions in Matlab.

### Intersection of the HNC-h-DEGs with Oct4, Sox2 and Nanog interactomes

The Oct4 interactome was built as the union of the Oct4 interactomes from Pardo *et al.* (2010), van den Berg *et al.* (2010), Ding *et al.* (2012) and Esch *et al.* (2013). The Sox2 interactome was taken from Fang *et al.* (2011), and the Nanog interactome from Gagliardi *et al.* (2013).

### Finding *in vitro* counterparts of the E3.25-E4.5 inner cells in the GEO database

We downloaded all 665 ESC transcriptomics profiles from the Affymetrix Mouse Genome 430 2.0 Array platform from the GEO database available at the time and RMA normalized it together with the *in vivo* data from Table 1. We averaged the single-cell transcriptomics profiles of each stage and differentiated cell type PE and EPI where applicable (E3.25-LNC, E3.25-HNC, E3.5PE, E3.5EPI, E4.5PE, E4.5EPI) and calculated the distance (using the Spearman correlation metric) to each ESC profile. From each of the six individual stage (and cell type) comparisons, we selected the ten closest ESC profiles. Next, we took the union of the six groups and we visualized the results in a heatmap.

### ScRNA-seq data processing

The SRA database for GSE84892, SRP079965, contains 292 single cell RNA-seq data from 16, 32 and 64 cell stage mouse embryos, out of which 122 were classified by as ICM by Posfai *et al.* (2017).

The single end RNA-seq data were produced on Illumina HiSeq 2000 platform. We aligned raw data to the GRCm38 genome using Tophat, and calculated the Fragments Per Kilobase of transcript per Million mapped reads (FPKM) using Cufflinks and in house software in Python 2.7. Variance stabilization was performed using log_2_ scaling. We selected the top high 80 DEGs upregulated in the late 32-cell ICM cells in relation to the early 32-cell ICM cells corresponding to a fold change 2.2.

## Acknowledgments

DG and MJ A-B. have been supported by grants DFG113/18 from Diputaci6n Foral de Gipuzkoa, Spain, Ministry of Economy and Competitiveness, Spain, MINECO grant BFU2016-77987-P and Instituto de Salud Carlos III (AC17/00012) co-funded by the European Union (Eracosysmed/H2020 Grant Agreement No. 643271).

